# Therapeutic potential of rWnt5A in curbing *Leishmania donovani* infection

**DOI:** 10.1101/2022.09.26.509479

**Authors:** Shreyasi Maity, Malini Sen

## Abstract

In this study, we demonstrated that administration of recombinant Wnt5A (rWnt5A) to *Leishmania donovani* (*L. donovani*) infected RAW 264.7 macrophages cause a significant reduction of infection that is associated with an increase in both cellular reactive oxygen species (ROS) and secreted IFN-γ/IL-10 ratio. Furthermore, rWnt5A administration appreciably blocks the progression of *L. donovani* infection established in a mouse model. This line of defense *in vivo* is associated with an elevation in immune cell associated ROS, and an activated T cell profile marked by increased IFN-γ and Granzyme B expression. As a corollary, infection associated disruption of splenic white pulp organization is markedly restrained. In summary, this study unveils the therapeutic potential of rWnt5A in curbing *L. donovani* infection and the progression of experimental visceral leishmaniasis through immune cell signaling. Additionally, it opens up new avenues of investigation into the use of rWnt5A as a therapeutic agent for restraining progression of the drug resistant refractory disease.

## Introduction

Visceral leishmaniasis (VL) or Kala-azar, caused by infection with the parasite *L. donovan*i turns out fatal if not appropriately treated. While several other drugs such as Amphotericin B and Miltefosine are now used in addition to the traditional antimony drugs to treat VL, the disease still persists in isolated pockets, especially in the rural areas (1–3). Although not endemic, refractory disease arising from infection with the drug resistant *L. donovani* strains remains a formidable challenge (4–6). The situation gets additionally complicated with the resurgence of the disease as Post Kala-azar Dermal Leishmaniasis (PKDL), although debate continues as to whether PKDL is a form of recurrent VL or a new disease (7–10). In this scenario it is important to have a clear concept of powerful immunomodulators that can potentially confront and curb infection. One immunomodulator that may be considered in this context is Wnt5A.

Wnt5A belongs to the Wnt family of secreted glycoprotein ligands that initiate signaling through binding to the Frizzled and/or ROR family receptors (11–14). While signaling by some Wnts, (canonical), for example Wnt3A categorically leads to the nuclear translocation of β-catenin, signaling by others (non-canonical), of which Wnt5A is a representative does not always involve β-catenin translocation. Despite such classification there is some degree of overlap between the canonical and non-canonical modes of Wnt signaling on account of homology between the ligand receptor pairs and common signaling intermediates (15, 16). Nevertheless, there are distinct differences in the outcome of signaling by the representative Wnts, perhaps on account of differences in the level of activities of the signaling intermediates. For instance, while prior activation of Wnt5A signaling blocks *L. donovani* infection, Wnt3A signaling promotes it (17, 18). In line with this observation we also demonstrated that the Wnt5A-Actin axis of macrophages is instrumental in determining the fate of pathogens in comparison to non-pathogens (19, 20).

In this article, based on the antagonistic role of Wnt5A signaling towards *L. donovani* infection, we explored the therapeutic potential of rWnt5A in blocking the progression of *L. donovani* infection in a mouse model. Our results indicate that administration of rWnt5A to *L. donovani* infected mice is able to reduce the infection load significantly.

## Materials & Methods

### Reagents

All tissue culture reagents and antibodies used in experiments are enlisted in Table 1.

**Table 1.**

| Reagents | Company | Catalog Number |
| --- | --- | --- |
| Anti-mouse IFN-γ ELISA set | BD Biosciences | 555138 |
| Anti-mouse IL-10 ELISA set | BD Biosciences | 555252 |
| Brefeldin | Calbiochem | 203729-1MG |
| BSA | SRL | SRL-83803 |
| BV605-rat anti-mouse CD8 | BD Biosciences | 563152 |
| Cell tracker green | Invitrogen | C2925 |
| DCFDA | Calbiochem | 287810-100MG |
| DMEM High Glucose Medium | Invitrogen | 12800017 |
| Fetal Bovine Serum | Invitrogen | 10082147 |
| Filipin | Sigma | F4767 |
| Giemsa | Himedia | S011 |
| Histopaque solution | Sigma | 10771-100ml |
| L-Glutamine | Invitrogen | 25030081 |
| Medium 199 | Invitrogen | 11150-059 |
| Methanol | Finar | 40930LC250 |
| PE-CF594 rat anti-mouse IFN-γ | BD Biosciences | 562333 |
| PE-Cyanine 7-rat anti-mouse Granzyme B | eBioscience | 25-8898-82 |
| PE-Cyanine 7-rat anti-mouse Ly6C | eBioscience | 25-5932-82 |
| PenStrep | Invitrogen | 15140122 |
| PerCP-Cy5.5 rat anti-mouse CD4 | BD Biosciences | 550954 |
| PerCP-Cy5.5 rat anti-mouse Ly6G | BD Biosciences | 560602 |
| Recombinant Wnt5A | Millipore | GF146 |
| RPMI 1640 | Invitrogen | 31800-022 |
| TMB solution | Merck | CL07-1000MLCN |
| Tween-20 detergent | Merck | 655205-250ML |
| V500-Syrian hamster anti-mouse CD3 | BD Biosciences | 560771 |

### Cell maintenance

RAW264.7 cell line was purchased from ATCC (ATCC©T1B71TM). The cells were maintained under conventional cell culture conditions at 37°C and 5% CO_2_ in DMEM high glucose medium supplemented with 10% FBS, 2 mM L-glutamine, 100 U/ml penicillin, and 100 mg/ml streptomycin.

Peritoneal cells harvested from BALB/c mice were resuspended in RPMI 1640 medium supplemented with 10% FBS, 2 mM L-glutamine, 100 U/ml penicillin, and 100 mg/ml streptomycin and kept under standard cell culture conditions (37°C and 5% CO_2_) for further experiments.

### Animal and parasite maintenance

8–12 weeks old male or female BALB/c mice were maintained in IVCS (Individually Ventilated Caging System) of the CSIR-IICB institutional animal facility with required access to food and water. A 12-hour day/night cycle, and constant temperature were preserved for animal maintenance. The *L. donovani* WHO reference strain AG83 [MHOM/IN/1983/AG83], procured from CSIR-IICB was propagated through BALB/c mice for maintenance in large numbers. The promastigote form of the parasite was maintained in Medium 199 supplemented with 20% FBS, 2 mM L-glutamine, 100 U/ml penicillin, and 100 mg/ml streptomycin at 22°C in a B.O.D. shaker incubator.

### Evaluation of rWnt5A mediated blockade of *L. donovani* infection

RAW264.7 macrophages were plated for 12 hours and then infected with *L. donovani* AG83 at 5 MOI for 4 hours. After 3 washes with 1X PBS, cells were treated with 50ng/ml of rWnt5A or PBS (vehicle control). After 6 hours of rWnt5A treatment, the media was changed and the cells were kept for another 6 hours in freshly added media. Subsequently, the supernatant was collected and stored for ELISA. The cells were harvested for either FACS or estimation of parasite burden, following fixation with absolute methanol and then staining with Giemsa.

For mice experiments, BALB/c mice were infected with *L. donovani* AG83 promastigotes (10^8^) through the intravenous route. One group of infected mice was sacrificed to check the infection load 15 days post-infection. Other groups were treated with either 100 ng rWnt5A or PBS through the intravenous route on the 15^th^ day and 30^th^ day after infection. The treated mice were then sacrificed on the 45^th^ day after infection to check the infection load in the spleen and liver through the Giemsa-stained imprint smears.

### Enumeration of parasite burden

The parasite burden in RAW264.7 cells was counted as the number of internalized amastigotes per 100 macrophages, by taking at least 20 microscopic fields using a Leica DFC450c camera at 100X magnification in the bright field of a Leica microscope.

For mice experiments, spleen and liver imprints were taken on clean glass slides and then fixed with chilled absolute methanol. Following fixation, the smears were stained with Giemsa. The Giemsa-stained micrographs were recorded with a Leica DFC450c camera at 100X magnification in the bright field of a Leica microscope. Leishman Donovan Unit (LDU) calculations were performed using at least 20 microscopic fields, by counting the amastigotes per 1000 host nuclei, and multiplying those numbers by the gram weight of the relevant organ.

### Blood smear preparation

Blood was taken from the tail vein and spread on a glass slide using another glass slide. The slide was then allowed to air dry before being stained with Giemsa. By keeping the slide submerged in softly running tap water, the extra stain was removed. After drying, the slide was examined using a Leica microscope with a 100X objective. Leica DFC450c camera was used to take the micrographs.

### ELISA (cytokine estimation)

IL-10 and IFN-γ levels in the harvested culture media from *L. donovani* infected RAW264.7 cell culture were measured with a BD ELISA set (IL-10 and IFN-γ) in accordance with the manufacturer’s instructions (www.bdbiosciences.com) in order to estimate the change in secreted cytokines after application of rWnt5A. The capture antibody was coated into 96-well polystyrene ELISA plates, which were incubated at 4° C overnight. Before blocking the plates for an hour with 10% FBS in 1X PBS, the plates were washed with PBST 3 times. Following the addition of 100 μl of the harvested media, the plates were incubated with a biotinylated detection antibody and streptavidin HRP. After about 30 min, TMB (substrate for HRP) was added. The color reaction was terminated by stop solution (2N H_2_SO_4_) and reading was obtained at 450 nm using an ELISA plate reader.

### Flow cytometry (analysis of ROS and T cell activation)

Spleen tissues from treated and control mice were minced and passed through a cell strainer to obtain single-cell suspensions. Following resuspension in RPMI, the cells were plated and left to incubate overnight under standard cell culture conditions. During the last 4 hours, Brefeldin A (3μg/ml) was added. Cells were then harvested and fixed with 1% paraformaldehyde. Following 3 washes with 1X PBS, the splenocytes were permeabilized with 0.1% Tween containing 0.5% BSA in 1X PBS. Subsequently, the cells were stained for an hour at 4°C with intracellular markers (PECF594 rat anti-mouse-IFN-γ and PE-Cyanine 7 rat anti-mouse-Granzyme B) and surface markers (V500 syrian hamster anti-mouse-CD3, PerCP-Cy5.5 rat anti-mouse-CD4, and BV605 rat anti-mouse-CD8) and prepared for FACS.

Separately, blood was drawn from the facial vein of both treated and control mice using 3.8% sodium citrate as an anticoagulant and PBMCs were isolated from the blood using a conventional Histopaque gradient (https://www.infinity.inserm.fr/wp-content/uploads/2018/01/PBMC-isolation-and-cryopreservation). For ROS and surface marker staining in PBMCs the cells were first incubated with the cell permeable dye DCFDA (10μM) for 20 minutes at 37°C and then washed 3 times with 1X PBS before staining for 1 hour at 4°C with antibodies against the cell surface markers V500 syrian hamster anti-mouse-CD3, PE-Cyanine 7-rat anti-mouse Ly6C, and PerCP-Cy5.5 rat anti-mouse Ly6G.

### Histological scoring

Histological scoring was carried out in accordance with a previous study (17). Briefly, during microscopic examination of the Hematoxylin and Eosin stained mouse spleen tissue sections, four different criteria were used to assess the degree of splenic white pulp structural organization: organized (OD: distinct germinal center and marginal zone), slightly disorganized (SD: slight loss of differentiation of germinal center and marginal zone), moderately disorganized (MD: indistinct germinal center and marginal zone) and intensely disorganized (ID: barely discernible white pulp and red pulp area). The number of white pulps in each category was divided by the total number of countable white pulps (at least 10 fields per mouse), and the ratio was multiplied by 100 to project the percent organization in terms of OD, SD, MD, and ID.

### Confocal microscopy

To check the level of cellular cholesterol, peritoneal exudates taken from BALB/c mouse was plated for 6 hours in chambered slide and then treated with either rWnt5A (50ng/ml) or PBS for 6 hours. The cells were then infected for 4 hours with cell tracker green labelled *L. donovani* AG83 at 5 MOI. For uninfected wells media was changed. Cells were incubated for a further 6 hours after being washed three times in PBS after 4 hours. Fresh 4% paraformaldehyde was used to fix the cells for 10 minutes, followed by three PBS washes. To quench the paraformaldehyde the cells were then incubated with 1.5 mg glycine/ml PBS for 10 minutes followed by staining with Filipin (1:1000 dilution) for 2 hours. The cells were then mounted with 60% glycerol after being washed three times with PBS. A Leica TCS-SP8 microscope was used to examine the cells.

### Statistical Analysis

Graph Pad Prism 5 software was used to analyze data using both paired and unpaired t-tests as appropriate. Expression for scatter plots and bar graphs is mean ± SEM. Statistics were considered significant at p ≤ 0.05. Annotation of significance was as follows: ns: not significant, *p ≤ 0.05, **p ≤ 0.005, ***p ≤ 0.0005.

### Ethics Statement

The IICB Animal Ethics Committee approved the use of animals. The approved identification codes are IICB/AEC/Meeting/2016/Aug, IICB/AEC/Meeting/Aug/2018/4 and IICB/AEC/Meeting/Sept/2019.

## Results and Discussion

### rWnt5A administration has inhibitory effect on *L. donovani* infection load in macrophages

In order to evaluate the effect of rWnt5A on *L. donovani* infection load we initially treated *L. donovani* infected RAW264.7 macrophages with rWnt5A. As clearly depicted in **Figure 1A**, the number of amastigotes were much reduced after rWnt5A treatment as opposed to just PBS (vehicle control) treatment indicating that rWnt5A induces parasite death. In connection with rWnt5A induced parasite reduction there was increase in the level of cell associated ROS (**Figure 1B**), implying that parasite death could in part be due to increase in ROS (21, 22). Increased ROS in the rWnt5A treated infected cells was in compliance with prior results depicting increased ROS in rWnt5A treated macrophages (18). Moreover, the ratio of secreted IFN-γ to IL-10 was higher in the *L. donovani* infected macrophages treated with rWnt5A as opposed to the corresponding controls (**Figure 1C**). The gating strategy for ROS+ RAW264.7 macrophages is illustrated in **Figure 1D**. In view of the anti-parasitic effects of IFN-γ and the pro-parasitic effects of IL-10 (23–25), this observation reinforced the antagonistic effect of Wnt5A toward *L. donovani* infection. Thus, we became interested to evaluate the therapeutic potential of rWnt5A on *L. donovani* infection in a mouse model.

**Figure 1:**
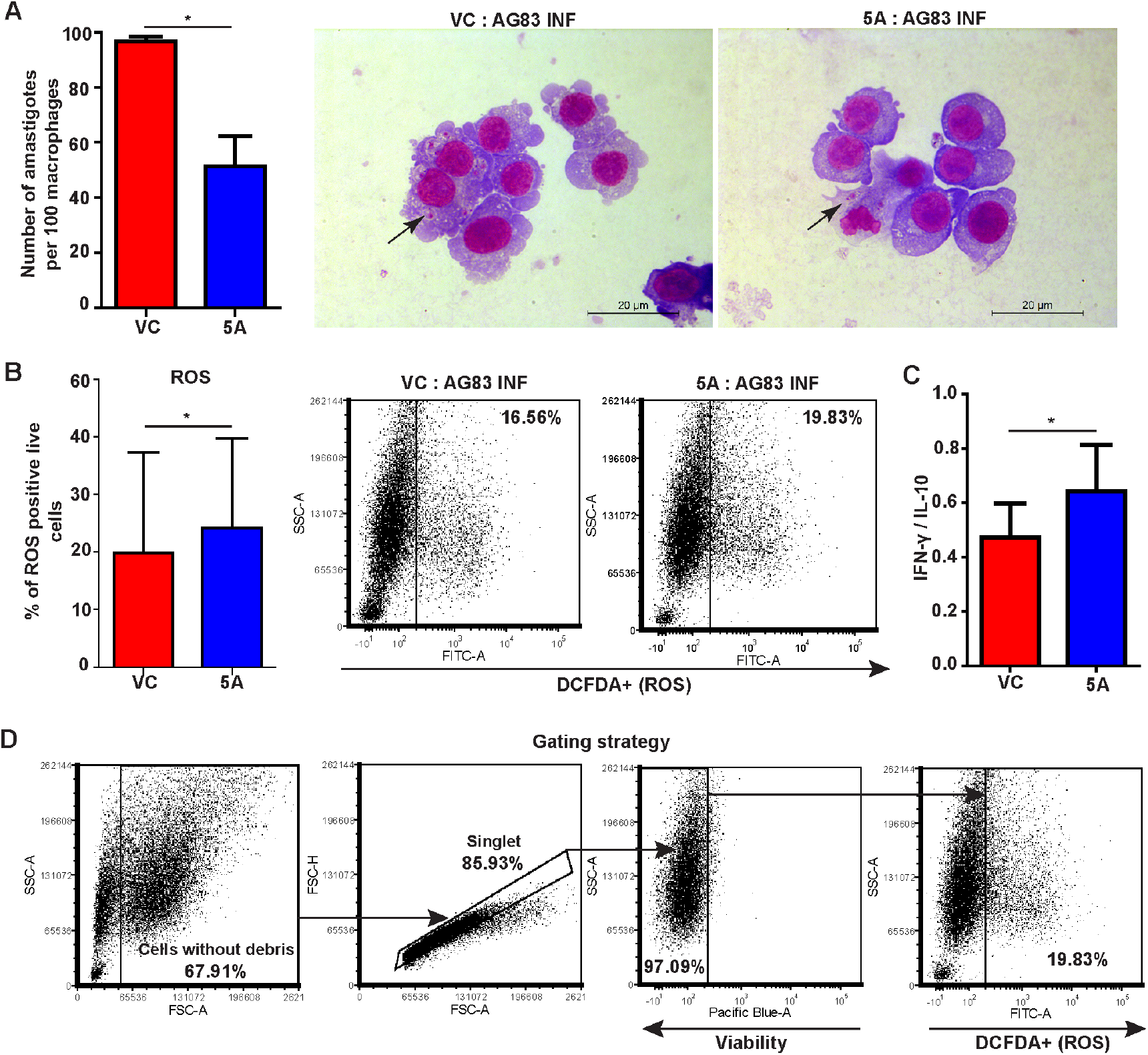
rWnt5A treatment causes restriction of *L. donovani* infection in RAW264.7 macrophages. **(A**) Bar graph and representative Giemsa-stained micrographs demonstrating decreased number of amastigotes (indicated by black arrows) in *L. donovani* infected RAW264.7 macrophages after rWnt5A (5A) treatment as compared to PBS:Vehicle Control (VC). (**B**) Bar graph and the representative FACS dot plots showing an increase in the level of cell-associated ROS in conjunction with reduction of parasites after rWnt5A treatment as compared to control. (**C**) Compared to the corresponding control, rWnt5A treatment of *L. donovani*-infected macrophages is linked with a higher ratio of secreted IFN-γ to IL-10. Paired t-test was used for statistical analysis. The following significance level was noted: *p ≤ 0.05. n=3 (number of experiments).

### rWnt5A induced reduction in *L. donovani* infection in a mouse model is associated with increase in cellular ROS and T cell activation

In order to test the potential of rWnt5A in curbing *L. donovani* infection, rWnt5A (100 ng) or PBS (vehicle control) was administered intravenously to *L. donovani* infected mice (10^8^ parasites/ mouse) in two doses; the first one on the 15th day after infection, and the second after 30 days. All experimental and control mice were subsequently sacrificed 45 days after infection following which Leishman Donovan Units (LDU) were enumerated separately from the liver and spleen of each mouse. That infection was established in mice within the 15 days before rWnt5A treatment was assessed initially by LDU enumeration in separate experiments (**Figure 2A**). As depicted in **Figure 2B**, both spleen and liver LDU were much lower in the infected mice that were treated with rWnt5A instead of just the vehicle control, confirming that rWnt5A administration leads to parasite reduction. The procedure of LDU enumeration is explained in the methods section.

**Figure 2:**
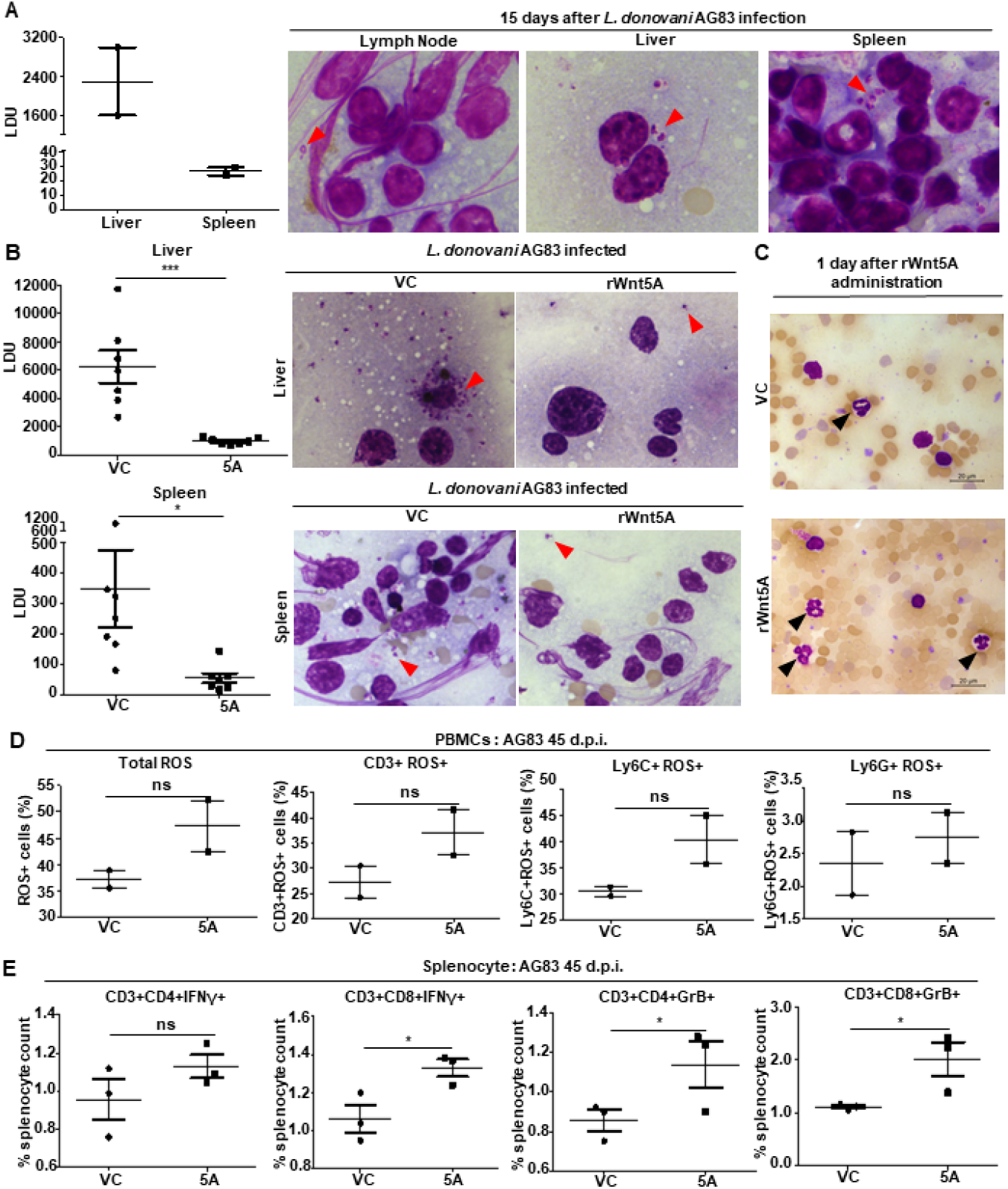
Increased cellular ROS and T cell activation are linked with rWnt5A-mediated suppression of *L. donovani* infection in a mouse model. (**A**) Graphical representation of LDU count and the Giemsa-stained imprint smears of liver, spleen and lymph node demonstrating parasite load after 15 days of infection. (**B**) Graphs of spleen and liver LDU count and representative Giemsa-stained imprint smears revealing that rWnt5A (5A) administration after infection causes a significant reduction in the infection load as compared to the vehicle control (VC). (**C**) Giemsa-stained images of blood smears showing an increase in the number of circulating neutrophils 1 day after rWnt5A treatment in *L. donovani* infected mice. (**D**) Graphs showing increased percentage of total ROS positive cells, CD3+ROS+ cells, Ly6C+ROS+ cells and Ly6G+ROS+ cells after rWnt5A treatment as compared to the vehicle control. (**E**) Graphs demonstrating that the *L. donovani*:AG83-infected rWnt5A-treated mice have higher percentage of IFN-γ and Gr-B+, CD3+CD4+ and CD3+CD8+ T cells compared to the vehicle control group. Unpaired t-test was used for statistical analysis. The following significance levels were noted: *p ≤ 0.05, ***p ≤ 0.0005, ns=not significant. n=2 to 7 (number of animals per group). dpi: days post infection.

The increase in circulating neutrophils in the infected mice one day after rWnt5A administration as opposed to PBS (vehicle control) administration as depicted by Giemsa staining of blood smears (**Figure 2C**) could at least in part be accountable for the rWnt5A induced parasite clearance. Assessment of cell associated ROS level in the circulating PBMC (monocytes, neutrophils and T cells) of the infected mice after rWnt5A or PBS administration revealed increase in the cell associated ROS of rWnt5A treated mice as compared to the corresponding controls (**Figure 2D**). The gating strategy for ROS+ PBMC is illustrated in **Supplementary Figure 1**. While the Wnt5A induced increase in the ROS level of monocytes and neutrophils, linked with parasite clearance, could be on account of phagosome associated NADPH oxidase activity, surge in T cell associated ROS could be mediated by the presentation of infected cell associated antigens to the T cells by antigen presenting cells (monocytes/macrophages). Furthermore, the accumulated ROS in T cells could cause transcriptional activation of immune defense mediators such as IFN-γ and GRB, which are linked with eradication of infection (26). Such activation profiles are likely to be reflected in the T cell repertoire of the spleens of rWnt5A treated *L. donovani* infected mice where LDU are much reduced as compared to the corresponding controls. Decrease in parasite load by rWnt5A in fact correlated with relatively higher levels of splenic CD4+ and CD8+ T cell associated IFN-γ and GRB, when compared to the corresponding controls (**Figure 2E**), corroborating that parasite clearance is linked with T cell activation. Activated T cells possibly prevent parasite persistence either through targeting and killing infected cells and/or activation of phagocytes through cytokine secretion. The gating strategy for T-cells is illustrated in Supplementary **Figure 2**.

We furthermore evaluated the spleen pathology of all infected mice, both rWnt5A and PBS administered, through histology in order to correlate parasite prevalence with disease status. In compliance with rWnt5A mediated parasite clearance, the spleens of the rWnt5A treated infected mice exhibited either minimal or unidentifiable disease states as evident from their near intact germinal centers/white pulp in contrast to the severe disease states of the infected controls as indicated by their highly unorganized splenic white pulp (**Figure 3A**) (27, 28). Histological scoring of spleen sections (explained in the methods section) confirmed significantly severe disease states in the control mice as opposed to the rWnt5A administered mice as depicted in **Figure 3B**.

**Figure 3:**
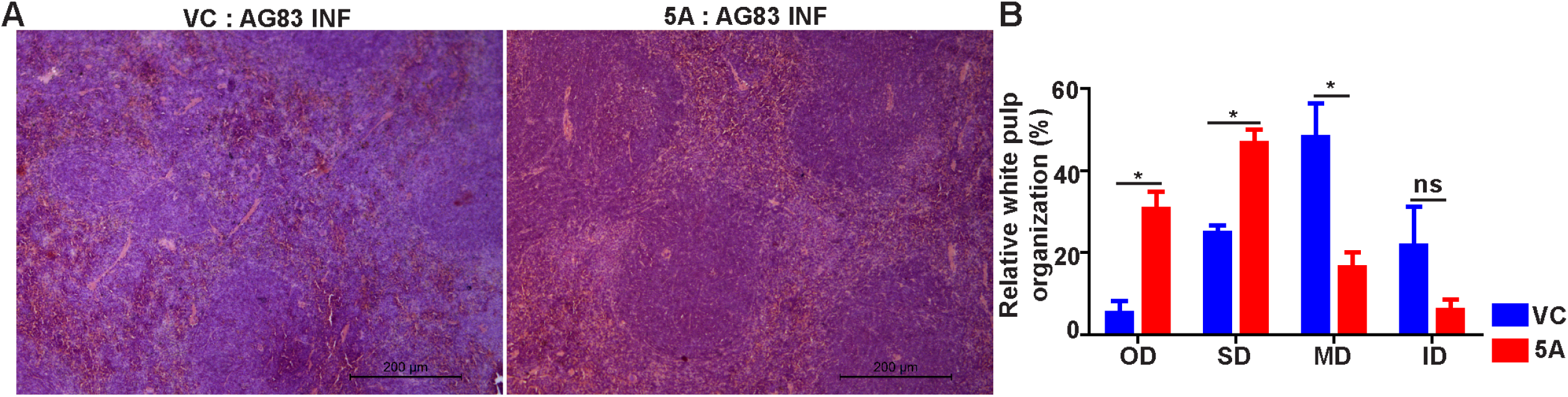
Comparative evaluation of spleen histopathology in rWnt5A vs PBS (VC) treated and *L. donovani* infected mouse. (**A**) Representative micrographs of H & E-stained spleen tissue sections (under 10X objective) collected from infected mice, which are either rWnt5A treated or PBS (VC) treated. (**B**) Graphical illustration showing various degrees of splenic white pulp disorganization in rWnt5A treated mice after 45 days of infection as compared to the vehicle control. Unpaired t-test was used for statistical analysis. The following significance levels were noted: *p ≤ 0.05, ns=not significant. n=2 (number of animals per group).

Taken together, our results reveal an as yet uncharacterized therapeutic role of rWnt5A in curbing *L. donovani* infection in a mouse model through the activation of monocytes/macrophages, neutrophils and T cells as outlined in a diagram (**Figure 4**). A death trigger of parasites in infected monocytes/macrophages is likely to be initiated at least partly by rWnt5A mediated modulations of the actin cytoskeleton, which is used by the parasites for building their niche (18). rWnt5A may also antagonize the spread of *L. donovani* infection through depletion of cholesterol from parasite niches on account of its potential in facilitating cholesterol egress (29). Our preliminary observation of cholesterol depletion in rWnt5A treated macrophages (both infected and uninfected) by confocal microscopy of Filipin stained cells is in compliance with such a concept. Reduction in *L. donovani* infection load by rWnt5A treatment was confirmed by staing the parasites with cell tracker green (**Figure S3**). Previously, in *L. donovani* infected mice pretreated with rWnt5A we had demonstrated the prophylactic ability of rWnt5A in preventing progression of *L. donovani* infection (17). The current demonstration of the possibility of rWnt5A mediated therapy for *L. donovani* infection bolsters the status of rWnt5A as an important player in the immune defense against infection with *L. donovani* and similar parasites. Most importantly, this study paves the way for new investigations into the use of rWnt5A as a therapeutic agent for curbing the progression of drug resistant VL.

**Figure 4:**
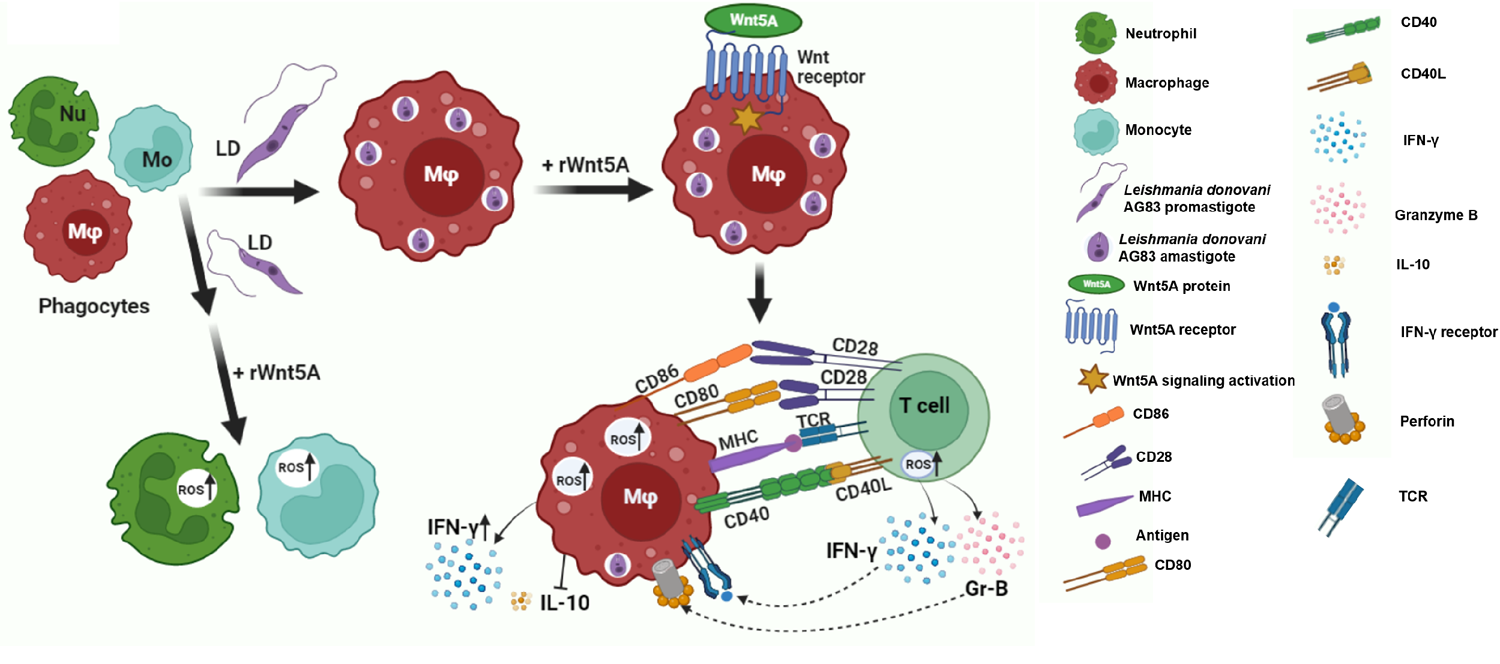
Model illustrates the therapeutic potential of rWnt5A in reducing *L. donovani* infection. The promastigote form of *L. donovani* (AG83) parasite infects the host phagocytes (mainly macrophages), where promastigote — amastigote conversion occurs. Macrophages carry the putative receptors for Wnt proteins. After administration of rWnt5A, signaling is triggered leading to ROS induction in monocytes/macrophages and T cells. T cells may get activated through antigen presentation. There may be interactions involving different costimulatory molecules: CD40 (present on macrophages)–CD40L (present on T-cells), CD80 (present on macrophages)-CD28 (present on T-cells), or CD86 (present on macrophages)-CD28 (present on T-cells). The activated T cells secrete IFN-γ and Gr-B. Simultaneous IFN-γ signaling pathway in macrophages can induce increased secretion of IFN-γ and decreased secretion of IL-10. With the aid of perforin, Gr-B maintains its capacity to kill infected cells. Increased production of ROS and activation of T-cells are associated with the therapeutic potential of rWnt5A.

## Acknowledgement

The authors acknowledge Tanmoy Dalui for FACS, Sounak Bhattacharya for confocal microscopy, Chandan Bhattacharya for tissue histology, CSIR-IICB for animal breeding and maintenance, Dr. Partha Chakraborty for the instrumental support and Soham Sengupta for assistance in cell culture.

---

**Supplementary Figure 1:**
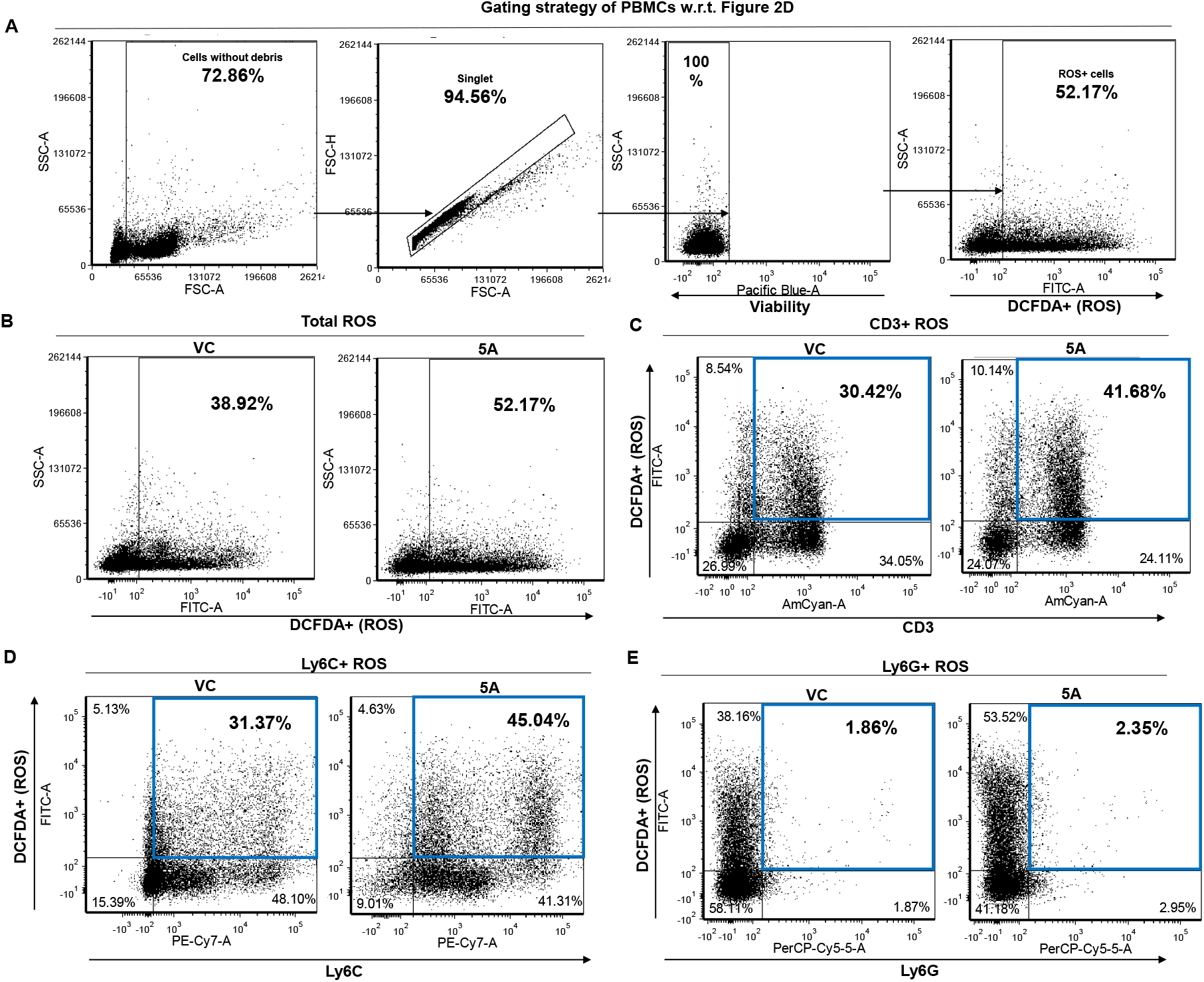
rWnt5A induced ROS production by PBMCs. **(A)** Depiction of gating strategy to identify total ROS+ PBMCs. **(B)** Examples of FACS dot plots showing an increase in the percentage of total ROS+ PBMCs collected from *L. donovani* infected mice after rWnt5A treatment as compared to the vehicle control. **(C-E)** Representative FACS dot plots showing an elevation in the percentage of CD3+ROS+ **(C)**, Ly6C+ROS+ **(D)** and Ly6G+ROS+ **(E)** PBMCs after rWnt5A treatment as compared to the vehicle control. The double positive cells are marked by a blue square on the upper right quadrant. The quadrant plots were made on the viable cells.

**Supplementary Figure 2:**
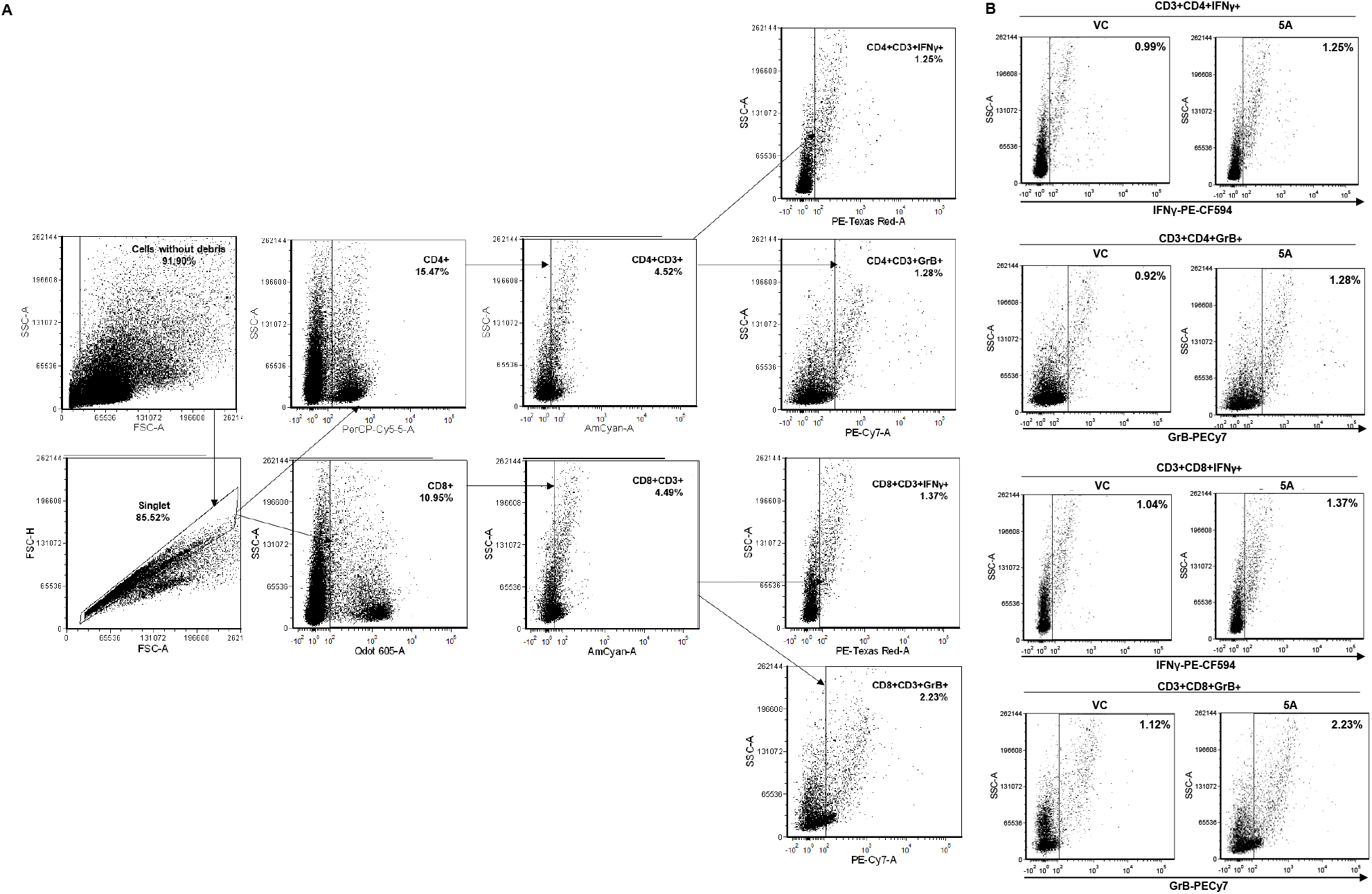
rWnt5A induced production of intracellular cytokines by splenic T cells. **(A)** Illustration of the gating method that was used to select out IFN-γ and Granzyme-B-positive CD3+CD4+ and CD3+CD8+ T cells. **(B)** Representative dot plots of splenocytes taken from infected mice which are either rWnt5Atreated or PBS treated, expressing IFN-γ and Granzyme-B positive CD3+CD4+ and CD3+CD8+ T cells.

**Supplementary Figure 3:**
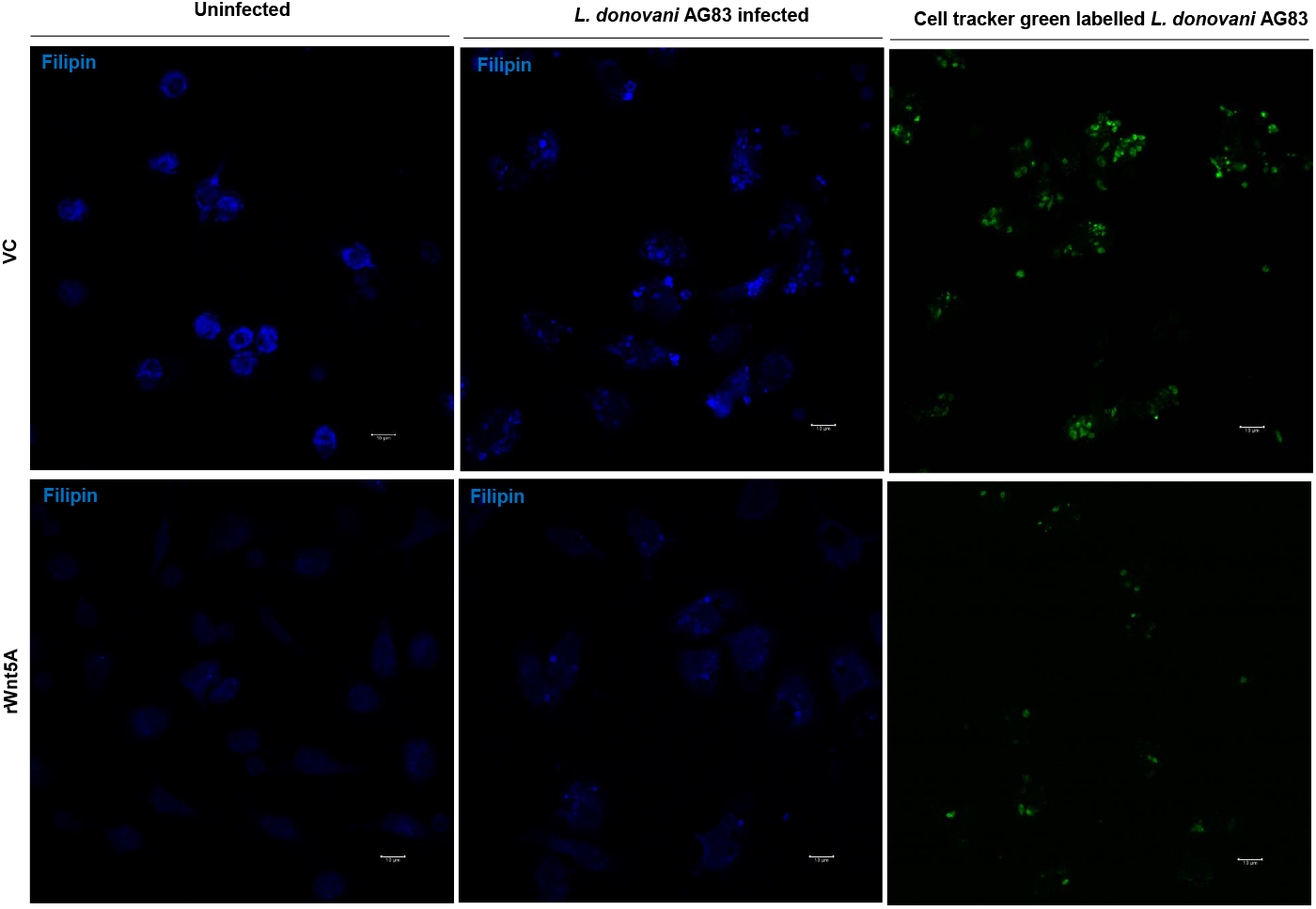
The restriction of *Leishmania donovani* infection caused by Wnt5A is connected to alteration of the cellular cholesterol. Confocal imaging of filipin-stained mouse peritoneal macrophages reveals a decrease in the amount of filipin-bound cholesterol in rWnt5A treated uninfected and rWnt5A treated *L. donovani* AG83 infected peritoneal macrophages as compared to the vehicle control (PBS). Cell tracker green labeled *L. donovani* AG83 parasites are quite low in number after rWnt5A treatment as compared to the vehicle control.

## References

1. Belbase, K., S. Shah, K. Dahal, S. B. Tiwari, and S. Neupane. 2022. Visceral leishmaniasis in non-endemic rural hilly region of Nepal: A case report. Clinical Case Reports 10: e05817.

2. Guan, Z., C. Chen, C. Huang, H. Zhang, Y. Zhou, Y. Zhou, J. Wu, Z. Zhou, S. Yang, and L. Li. 2021. Epidemiological features and spatial–temporal distribution of visceral leishmaniasis in mainland China: a population-based surveillance study from 2004 to 2019. Parasites & Vectors 14: 517.

3. Guha, S., M. Chatterjee, P. Saha, S. Ganguly, A. Chakraborty, N. Ganguly, A. Maji, and S. Das. 2017. EPIDEMIOLOGICAL STUDIES ON LEISHMANIASIS IN MURSHIDABAD DISTRICT OF WEST BENGAL, INDIA. International Journal of Recent Scientific Research 8: 1660–21664.

4. Chappuis, F., S. Sundar, A. Hailu, H. Ghalib, S. Rijal, R. W. Peeling, J. Alvar, and M. Boelaert. 2007. Visceral leishmaniasis: what are the needs for diagnosis, treatment and control? Nat Rev Microbiol 5: 873–882.

5. Alves, F., G. Bilbe, S. Blesson, V. Goyal, S. Monnerat, C. Mowbray, G. Muthoni Ouattara, B. Pécoul, S. Rijal, J. Rode, A. Solomos, N. Strub-Wourgaft, M. Wasunna, S. Wells, E. E. Zijlstra, B. Arana, and J. Alvar. 2018. Recent Development of Visceral Leishmaniasis Treatments: Successes, Pitfalls, and Perspectives. Clin Microbiol Rev 31: e00048–18.

6. Ponte-Sucre, A., F. Gamarro, J.-C. Dujardin, M. P. Barrett, R. López-Vélez, R. García-Hernández, A. W. Pountain, R. Mwenechanya, and B. Papadopoulou. 2017. Drug resistance and treatment failure in leishmaniasis: A 21st century challenge. PLOS Neglected Tropical Diseases 11: e0006052.

7. Gedda, M. R., B. Singh, D. Kumar, A. K. Singh, P. Madhukar, S. Upadhyay, O. P. Singh, and S. Sundar. 2020. Post kala-azar dermal leishmaniasis: A threat to elimination program. PLOS Neglected Tropical Diseases 14: e0008221.

8. Mukhopadhyay, D., J. E. Dalton, P. M. Kaye, and M. Chatterjee. 2014. Post kala-azar dermal leishmaniasis: an unresolved mystery. Trends Parasitol 30: 65–74.

9. Zijlstra, E. E. 2019. Biomarkers in Post-kala-azar Dermal Leishmaniasis. Frontiers in Cellular and Infection Microbiology 9.

10. Zijlstra, E. E., A. M. Musa, E. a. G. Khalil, I. M. el-Hassan, and A. M. el-Hassan. 2003. Post-kala-azar dermal leishmaniasis. Lancet Infect Dis 3: 87–98.

11. Minami, Y., I. Oishi, M. Endo, and M. Nishita. 2010. Ror-family receptor tyrosine kinases in noncanonical Wnt signaling: Their implications in developmental morphogenesis and human diseases. Developmental Dynamics 239: 1–15.

12. Schulte, G., and V. Bryja. 2007. The Frizzled family of unconventional G-protein-coupled receptors. Trends in Pharmacological Sciences 28: 518–525.

13. Nishita, M., S. Itsukushima, A. Nomachi, M. Endo, Z. Wang, D. Inaba, S. Qiao, S. Takada, A. Kikuchi, and Y. Minami. 2010. Ror2/Frizzled Complex Mediates Wnt5a-Induced AP-1 Activation by Regulating Dishevelled Polymerization. Molecular and Cellular Biology 30: 3610–3619.

14. Wnt signaling: a common theme in animal development.

15. Veeman, M. T., J. D. Axelrod, and R. T. Moon. 2003. A Second Canon: Functions and Mechanisms of β-Catenin-Independent Wnt Signaling. Developmental Cell 5: 367–377.

16. Mikels, A. J., and R. Nusse. 2006. Purified Wnt5a Protein Activates or Inhibits β-Catenin–TCF Signaling Depending on Receptor Context. PLOS Biology 4: e115.

17. Maity, S., A. Chakraborty, S. K. Mahata, S. Roy, A. K. Das, and M. Sen. 2022. Wnt5A Signaling Blocks Progression of Experimental Visceral Leishmaniasis. Front Immunol 13: 818266.

18. Chakraborty, A., S. P. Kurati, S. K. Mahata, S. Sundar, S. Roy, and M. Sen. 2017. Wnt5a Signaling Promotes Host Defense against Leishmania donovani Infection. The Journal of Immunology 199: 992–1002.

19. Jati, S., S. Kundu, A. Chakraborty, S. K. Mahata, V. Nizet, and M. Sen. 2018. Wnt5A Signaling Promotes Defense Against Bacterial Pathogens by Activating a Host Autophagy Circuit. Front. Immunol. 9: 679.

20. Jati, S., S. Sengupta, and M. Sen. 2021. Wnt5A-Mediated Actin Organization Regulates Host Response to Bacterial Pathogens and Non-Pathogens. Frontiers in Immunology 11: 3803.

21. Basu, M., and P. K. Das. 2019. Role of Reactive Oxygen Species in Infection by the Intracellular Leishmania Parasites. In Oxidative Stress in Microbial Diseases S. Chakraborti, T. Chakraborti, D. Chattopadhyay, and C. Shaha, eds. Springer, Singapore. 297–311.

22. Murray, H. W., and C. F. Nathan. 1999. Macrophage Microbicidal Mechanisms In Vivo: Reactive Nitrogen versus Oxygen Intermediates in the Killing of Intracellular Visceral Leishmania donovani. J Exp Med 189: 741–746.

23. Murphy, M. L., U. Wille, E. N. Villegas, C. A. Hunter, and J. P. Farrell. 2001. IL-10 mediates susceptibility to Leishmania donovani infection. European Journal of Immunology 31: 2848–2856.

24. Murray, H. W., C. M. Lu, S. Mauze, S. Freeman, A. L. Moreira, G. Kaplan, and R. L. Coffman. 2002. Interleukin-10 (IL-10) in Experimental Visceral Leishmaniasis and IL-10 Receptor Blockade as Immunotherapy. Infect Immun 70: 6284–6293.

25. Tsagozis, P., E. Karagouni, and E. Dotsika. 2005. Function of CD8+ T lymphocytes in a self-curing mouse model of visceral leishmaniasis. Parasitology International 54: 139–146.

26. Yarosz, E. L., and C.-H. Chang. 2018. The Role of Reactive Oxygen Species in Regulating T Cell-mediated Immunity and Disease. Immune Netw 18: e14.

27. Hermida, M. d’El-Rei, C. V. B. de Melo, I. D. S. Lima, G. G. de S. Oliveira, and W. L. C. Dos-Santos. 2018. Histological Disorganization of Spleen Compartments and Severe Visceral Leishmaniasis. Front Cell Infect Microbiol 8: 394.

28. Silva-O’Hare, J., I. S. de Oliveira, T. Klevorn, V. A. Almeida, G. G. S. Oliveira, A. M. Atta, L. A. R. de Freitas, and W. L. C. dos-Santos. 2016. Disruption of Splenic Lymphoid Tissue and Plasmacytosis in Canine Visceral Leishmaniasis: Changes in Homing and Survival of Plasma Cells. PLOS ONE 11: e0156733.

29. Awan, S., M. Lambert, A. Imtiaz, F. Alpy, C. Tomasetto, M. Oulad-Abdelghani, C. Schaeffer, C. Moritz, D. Julien-David, S. Najib, L. O. Martinez, R. L. Matz, X. Collet, R. Silva-Rojas, J. Böhm, J. Herz, J. Terrand, and P. Boucher. 2022. Wnt5a Promotes Lysosomal Cholesterol Egress and Protects Against Atherosclerosis. Circ Res 130: 184–199.

